# Effect of aging and *Varroa* parasitism on the paracellular and transcellular permeability of the honeybee blood-brain barrier

**DOI:** 10.1101/2024.09.27.615259

**Authors:** Tyler Quigley, Gro Amdam

## Abstract

Honeybees (*Apis mellifera*) provide crucial pollination services to agricultural systems globally, however, their healthspan in these contexts is constantly at risk. Agricultural environments impose a variety of sublethal stressors onto honeybees, including parasites, pathogens, pesticides, and poor nutrition. Synergies between age, age-associated tasks, and these stressors are believed to underlie colony failure trends of the past decade. Identifying the mechanisms by which age and stressors impact honeybee physiology is an important priority in protecting honeybee and other pollinator populations. An underexplored physiological structure in honeybees is the blood-brain barrier, a protective layer of cells that surround the brain. Here, we assess a key dimensions of blood-brain barrier function, paracellular and transcellular permeability to molecules in the hemolymph. We assess these modes of permeability in multiple age groups and after exposure to varying levels of infestation by the parasitic mite *Varroa destructor* during development. Our results demonstrate that the paracellular permeability of the honeybee blood-brain barrier is stable across their lifespan and upon *Varroa* exposure. In contrast, we found that transcellular permeability is increased in honeybees exposed to a high *Varroa* load. These results demonstrate how age and stress variably impact a primary protective structure of the honeybee central nervous system, which may lead to targeted interventions for protecting honeybee healthspan. The assay developed here may be easily applied to different aging- and stress contexts, further enabling studies focused on understanding maintenance and decline of the honeybee blood-brain barrier.

## Introduction

Over the past half century, agricultural intensification has become the primary driver of insect population decline worldwide.^1^ Paradoxically, modern agricultural practices impose a variety of risk factors onto wild and managed bee pollinators which are estimated to contribute hundreds of billions of USD to global crop production.^2–4^ These risk factors include pests, pathogens, pesticides, climate stress, poor nutrition, and poor management practices, each of which has distinct consequences for pollinator health.^5^ Pollinator health can be defined as “a state that allows individuals to live longer and/or reproduce more, even in the presence of pathogens, thus providing more ecological services.”^6^ Efforts to improve pollinator health are thus mutually beneficial between our species and the insect species we rely so heavily upon. Fortunately, remediating and protecting pollinator populations has become a priority for a variety of stakeholders, and recent policy reports have identified key priority areas to improve insect pollinator health specifically.^2,7,8^ Among these priorities is research to increase our understanding of the fundamental biological mechanisms by which stressors impact pollinator insect health.

Honeybees (*Apis mellifera*) are the most commonly managed insect pollinator species worldwide, and many crop production systems depend specifically on honeybee pollination.^9^ Nevertheless, annual honeybee colony losses have remained high across Europe, Canada, and the United States for over a decade.^10^ A “colony loss” can manifest in multiple ways, but the decline in colony performance over time leading up to a loss is a result of the collective decline in the health and performance of colony members, i.e. their ability to perform the tasks essential to colony survival across their lifespan.^5,11–13^

Understanding how risk factors impact honeybee healthspan can provide a path forward for improving individual and colony health. Although a risk factor may affect multiple physiological systems at once, a honeybee’s performance of a task is mediated by the central nervous system (CNS). Indeed, CNS dysfunction and decline seem to be a primary link between environmental stress, reduced individual healthspan, and colony failure.^11,14^ The neuroanatomy and cognitive architecture of the honeybee CNS is well understood as honeybees are a well-established neurobiological model useful for probing the fundamental principles that govern animal behavior, brain health, and neurological disease and aging.^15,16^ By leveraging and expanding this toolkit, we can better resolve the mechanisms by which risk factors dysregulate the CNS of honeybees over their life course, and contribute to reduced lifetime performance at individual and colony levels.

An underexplored region of the honeybee brain which may mediate the impact of stress on the honeybee CNS is the blood-brain barrier.^1^ The blood-brain barrier is a selective cellular barrier which regulates the access of molecules and signals from peripheral tissues to the brain.^17,18^ Its primary function is to maintain the strict ionic environment necessary for optimal neuronal signaling, however, it is also enriched in a variety of active transport systems responsible for the exchange of ions, nutritious metabolites, hormones xenobiotics, drugs, and metabolic waste products between the brain and hemolymph.^19^ The blood-brain barrier also acts as a homeostatic sensor which transduces information about the physiological state of an animal towards neuronal circuits.^20,21^ Despite differences in the cellular makeup and broad structure, many of the molecular systems that form and govern the blood-brain barrier are conserved between mammalian and insect blood-brain barriers.^22,23^ The homology of these systems provide a uniquely robust basis for applying comparative insights across model species, and has been particularly useful in understanding the mechanisms of aging-related decline in blood-brain barrier function.^24,25^

A key dimension of blood-brain barrier integrity is its permeability to molecules in circulation.^26^ In many other species, the permeability of the blood-brain barrier is dynamic across individual’s lifespan and can be impacted by factors including nutrition, environmental stress, disease, and aging.^27–29^ Although an animal’s behavior is coordinated by neurons, neuronal function can be severely impaired by a leaky blood-brain barrier.^17^ The consequences of increased blood-brain barrier permeability include increased oxidative stress, neurodegeneration, and cognitive decline.^30,31^ These physiological deficits are also observed in honeybees upon increasing age and exposure to environmental stress. It is plausible to suspect then that effects of age and stress on honeybee healthspan are mediated by changes in blood-brain barrier permeability. Understanding the mechanisms by which these effects impact honeybee healthspan can inform the development of novel interventions to improve honeybee health.

Here, we present an adapted insect blood-brain barrier permeability assay to measure the transcellular and paracellular permeability in honeybees.^32–34^ We applied this assay to honeybee workers to assess how blood-brain barrier permeability changes with age and how it is impacted by *Varroa destructor* parasitism. The assay described herein is simple and low-tech, which lends itself to adaptation for assessing insect blood-brain barrier permeability in diverse aging and stress contexts, thereby presenting new paths to understanding these impacts on a structure crucial for brain homeostasis.

## Methods

### Animals

Honeybees for this study were sourced from colonies were maintained at the Arizona State University Bee Lab in Mesa, Arizona. Collections occurred between Fall 2020 and Spring 2022.

### Sample Collection

#### Aging

To assess how honeybee blood-brain barrier permeability changes with age, we collected nurse bees (pre-foraging), young foragers (<14 days foraging), and old foragers (>14 days foraging). To collect these age groups, a brood frame was removed from a colony and placed in a wire cage in an incubator overnight at 33°C and 70% humidity. The next day, approximately 250 newly emerged bees were obtained from this frame, marked on the abdomen, and placed back into the hive (nurse bees). On the same day as the newly emerged bees were placed in the hive, approximately 1,500 foragers were marked with a different color upon their return from foraging (old foragers). Fourteen days later, another subset of 200 foragers were marked upon their return from foraging (young foragers). The day after young foragers were marked, five bees of each group were collected and brought into the lab for dye permeability assays.

#### Varroa

To assess how infection with *Varroa* impacts honeybee blood-brain barrier permeability, we collected recently emerged honeybees exposed to 0-1 mite and 2-4 mites during their development. Colonies with *Varroa* infestation were identified and left untreated for the duration of sampling each Fall or Spring season. To collect newly emerged honeybees, brood frames with actively emerging brood were collected and observed indoors under low light. As bees emerged, they were collected with soft forceps. Adult mites present on the honeybee and within its cell were counted and removed. Honeybees were paint marked with a color corresponding to the number of mites found on the bee and in the cell and placed into a wire mesh cylinder. Sampling occurred over multiple days with ∼10 total bees collected in each session. After each sampling session, the single wire mesh cage was placed into a host colony with no *Varroa* infestation. The specimens aged one day in the host colony to eliminate confounding development factors that may influence blood-brain barrier permeability. After 24 hours, the wire mesh cage was removed, brought into the lab for dye permeability assays.

### Dye permeability assay

We assessed paracellular and transcellular blood-brain barrier permeability in honeybees using an assay adapted from previous studies with honeybees and *Drosophila*.^32,35^ These modes of permeability are assessed with dyes which have distinct interactions with the blood-brain barrier. Paracellular permeability refers to the movement of molecules through the intercellular space between cells. Paracellular integrity is maintained at the blood-brain barrier by tight junctions between blood-brain barrier cells, minimizing the intercellular space through which molecules can pass to enter the brain.^19^ We used a Texas Red-conjugated 10 kDa dextran (TRD) (Invitrogen D-1863) to probe paracellular permeability, as this is the smallest molecular weight which is prevented from passing through the insect blood-brain barrier under healthy conditions.^32,34^ Transcellular permeability refers to the movement of molecules across the lipid bilayers of and through the cells that make up the blood-brain barrier.^19^ One way in which the blood-brain barrier regulates this pathway is by maintaining a high concentration of ATP-binding cassette (ABC) transporters on the apical membrane of blood-brain barrier cells.^36^ We used the dye Rhodamine B (Rho B) (Sigma R6626) to probe transcellular permeability, as it is permeable to cell membranes and is a substrate for the ubiquitous ABC transporter p-glycoprotein (P-gp), and possibly others.^35^

To perform the assay, specimens were secured into a custom mount and placed under a dissecting microscope. Using a surgical blade, the hair on the left side head was shaved down and a sliver of cuticle was removed from the head, exposing the tissue inside. Effort was made to minimize the size of the cut and the handling of the removed cuticle so as minimize disturbance to the underlying tissue. Often a layer of connective tissue remained immediately inside the incision; this gently teared to allow free dye permeation into the head capsule.

Either 1 uL of 0.125 mg/mL of RhoB dye or 1 uL of 5 mM 10-kDa dextran dye was then injected into the head capsule using a bevel-tip syringe (Lab Depot, 002105). Once injected, the bees were kept in the mounts, and placed onto a wet paper towel and covered with a cardboard box to reduce desiccation at the incision. After 45 minutes of exposure to the dye, bees were decapitated, and heads placed into a 1% paraformaldehyde solution. Brains were dissected under fixative, cleaned of surrounding tissue, and placed into a well in a 96 well plate. Wells were filled with 50 uL 0.1% SDS solution. SDS breaks up cell walls, releasing dye. Each brain was further processed within the well repeated pressing with a pestle for 30 seconds. Once all brains were dissected, placed, and processed in the well plate, the dye concentrations present in each well of the well plate was analyzed on a BioTek HT1 well plate reader (Excitation: 535 Emission: 595).

### Analysis

The relative fluorescence values derived from the brains of each age groups and each mite load group was used to compare paracellular and transcellular permeability between the groups. Comparisons of permeability among age groups were tested using ANOVA tests. Comparisons of permeability between mite load groups were tested using t-tests. Data is presented as boxplots depicting the median (bold center line), interquartile ranges (span of box), range within 1.5 times the interquartile range (whiskers) and outliers. All analyses were performed in R.

## Results

### Paracellular Permeability

We did not find a significance difference in dextran dye in the honeybee brains of nurses, young foragers, and old foragers increased with age (ANOVA, p=0.19) (Fig. 1A). Similarly, we did not find a significant difference in brain dextran concentration between honeybee workers who emerged with a low mite load versus a high mite load (t-test, p=0.424) (Fig. 1B).

**Figure 1:**
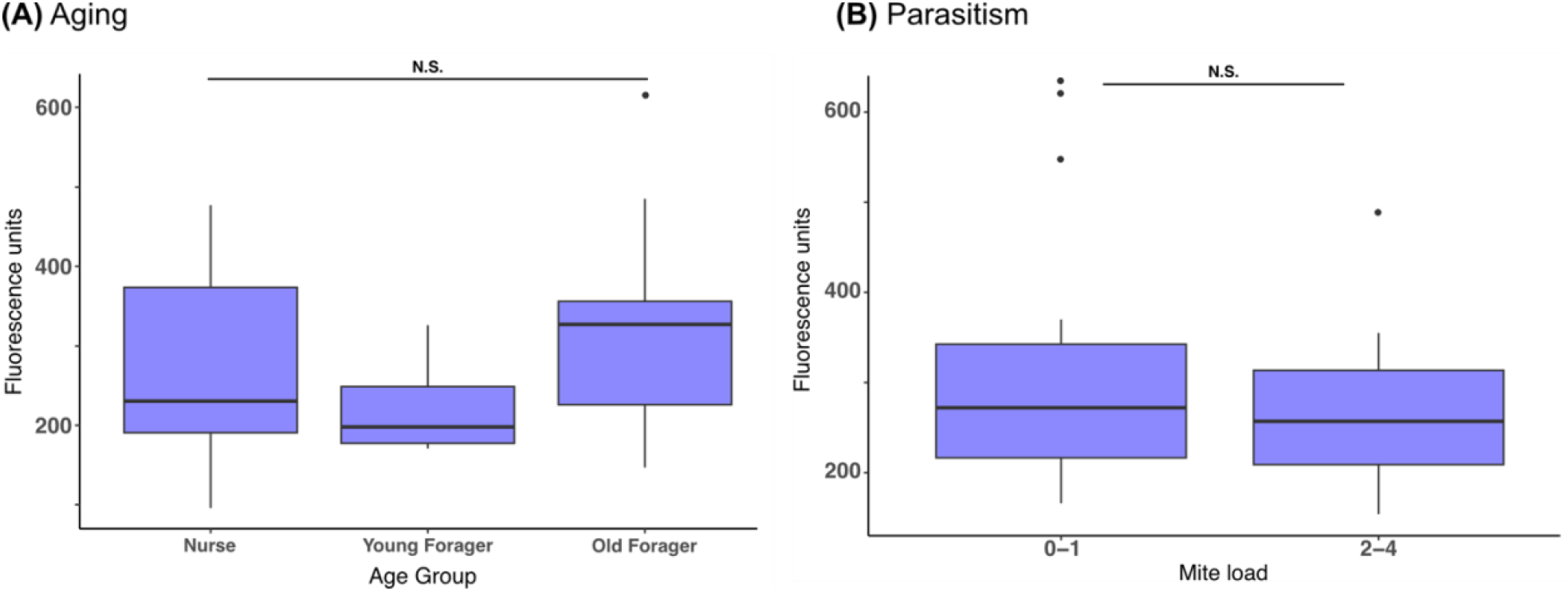
Paracellular permeability of the honeybee blood-brain barrier. Paracellular permeability of the honeybee blood-brain barrier. The amount of fluorescence emission from the brains of dextran-injected specimens is a quantitative measure of the paracellular permeability of the blood-brain barrier. There was no significant difference in dextran concentration amongst age groups (ANOVA, p=0.19) and mite load groups (t-test, p=0.424).

**Table 1:** Summary statistics for paracellular permeability data.

|  | n | Mean<br>fluorescence | df | F | *p* |
| --- | --- | --- | --- | --- | --- |
| Aging |  |  | ANOVA |  |  |
| Nurse | 8 | 270.1 | 2 | 1.799 | 0.19 |
| Young<br>Forager | 8 | 220.3 |  |  |  |
| Old Forager | 9 | 328.4 |  |  |  |
| Parasitism |  |  | t-test |  |  |
| 0-1 mite | 22 | 305.5 | 0.812 | 26.9 | 0.424 |
| 2-4 mite | 8 | 273.1 |  |  |  |

### Transcellular Permeability

The mean Rho B concentration in the brains of nurses, young foragers, and old foragers increased with age, however, the differences between groups were not significant (ANOVA, p = 0.21) (Fig. 2A). The brain Rho B concentration in honeybees that emerged with a high mite load was significantly higher than found in the brains of honeybees emerged with a low mite load (t(37.9) = -3.00, p = 0.0047***) (Fig. 2B).

**Figure 2:**
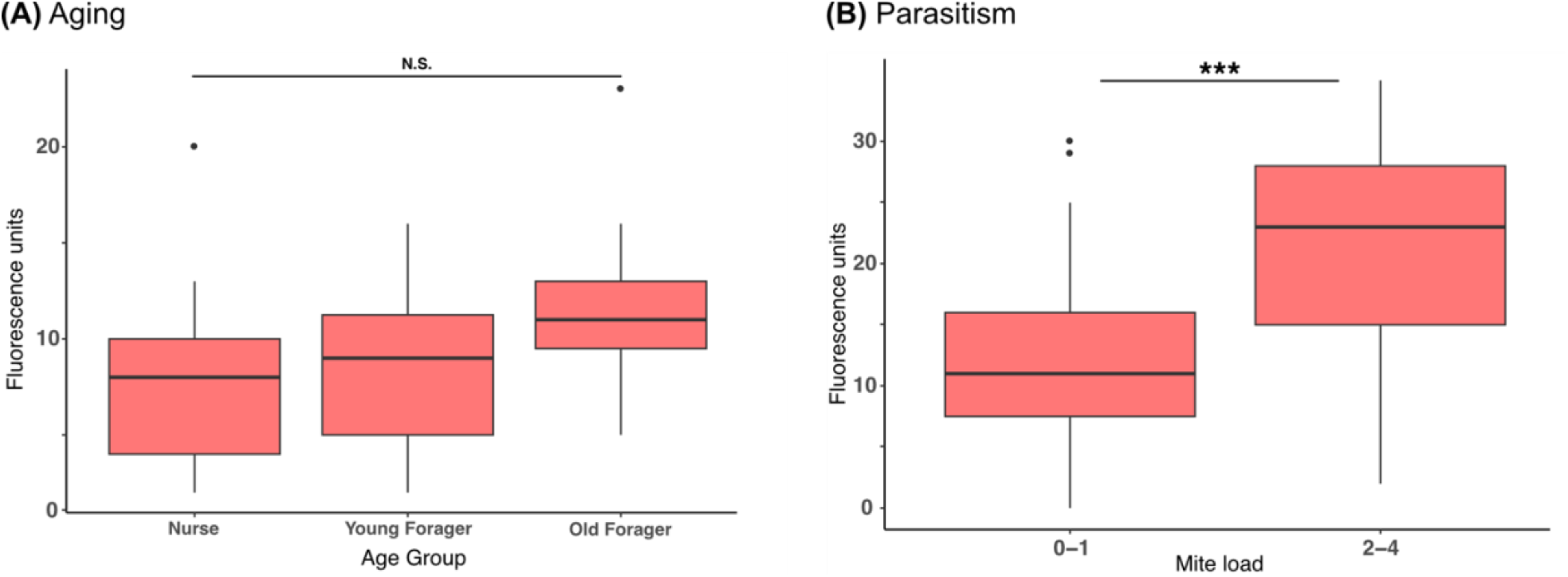
Transcellular permeability of the honeybee blood-brain barrier. Fluorescence signals from high and low mite load honeybee workers. (a) There was no difference in Rho B dye signal from the brains of these two groups (t(26.9) = 0.812, p = 0.424). (b) The Rho B signal observed from the brains of workers which emerged with two or more mites is significantly higher that the signal observed from the brains of workers which emerged with zero or one mite (t(37.2) = -3.02, p = 0.0045).

**Table 2:**
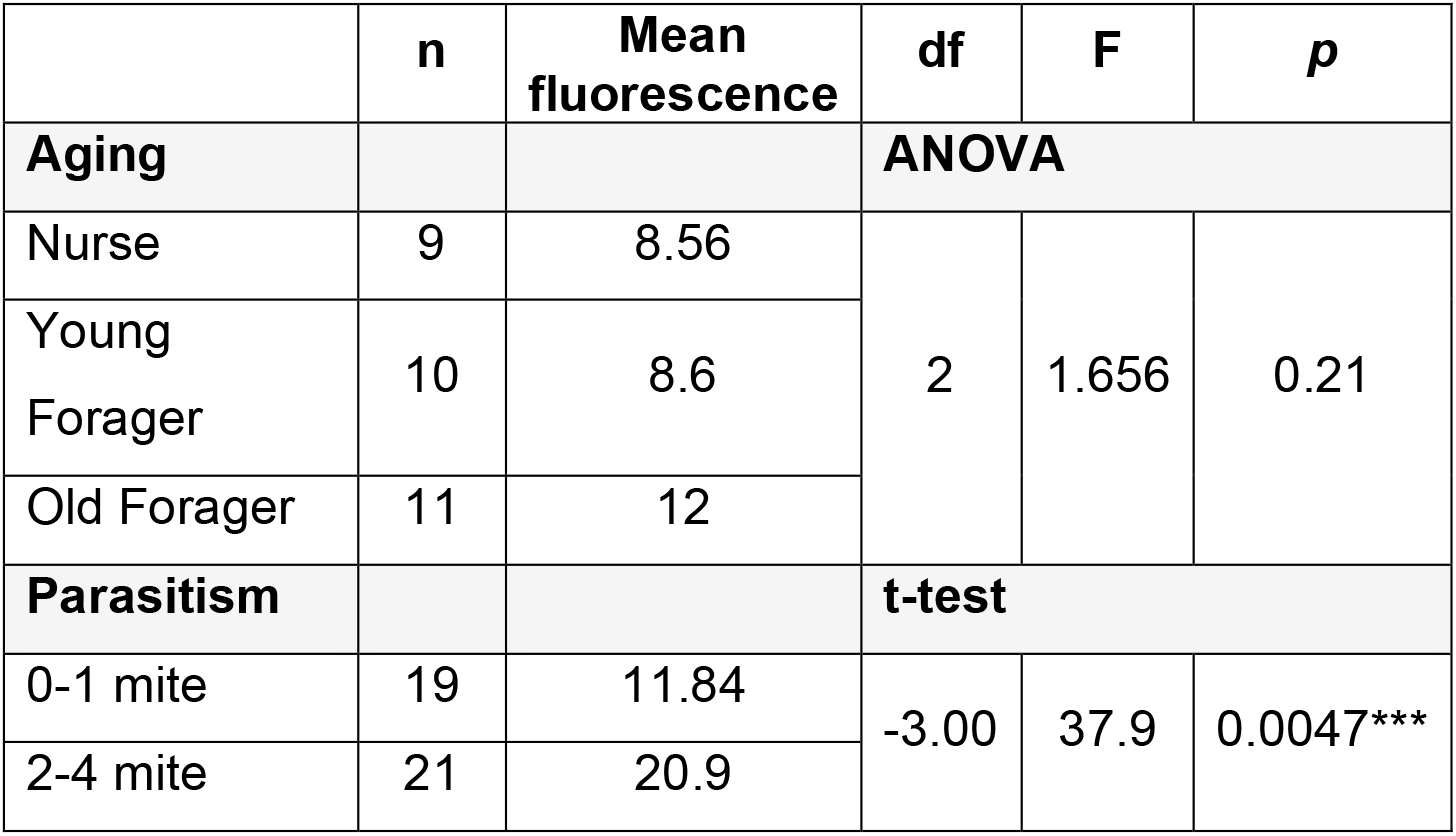
Summary statistics for transcellular permeability data.

|  | n | Mean<br>fluorescence | df | F | p |
| --- | --- | --- | --- | --- | --- |
| Aging |  |  | ANOVA |  |  |
| Nurse | 9 | 8.56 | 2 | 1.656 | 0.21 |
| Young<br>Forager | 10 | 8.6 |  |  |  |
| Old Forager | 11 | 12 |  |  |  |
| Parasitism |  |  | t-test |  |  |
| 0-1 mite | 19 | 11.84 | -3.00 | 37.9 | 0.0047*** |
| 2-4 mite | 21 | 20.9 |  |  |  |

## Discussion

The healthspan and performance of a honeybee worker is closely linked to their CNS health.^14,37^ The CNS’s primary line of defense against potentially harmful endogenous and exogenous factors is the blood-brain barrier. Here, we compared the paracellular and transcellular permeability of the honey blood-brain barrier across different worker age groups and between workers with low and high mite load. We did not find a difference in paracellular permeability across age groups or between mite load levels (Fig. 1). Conversely, we found a non-significant increase in transcellular permeability in workers with age (Fig. 2A), and significantly higher transcellular permeability in workers who emerged with a high mite load versus those emerged with a low mite load (Fig. 2B).

The insect blood-brain barrier is formed by the neural lamella and two layers of distinct glial subtypes.^38,39^ The neural lamella is an acellular fibrous matrix that sheaths the entire nervous system. Perineurial glia (PG) compose the most outer cellular layer and subperineurial glia (SPG) form the layer immediately beneath PG. Paracellular diffusion is primarily blocked by a dense network of junctional protein complexes that bind the maze-like interface between SPG cells.^40,41^ We assessed the viability of the paracellular barrier with a 10 kDa dextran dye, which is normally excluded by the SPG layer, but can permeate if the paracellular diffusion barrier is disrupted.^32^ This assay is often used to assess the impact of genetic disruptions to the *Drosophila* BBB, however it has also been employed to demonstrate how paracellular integrity is disrupted in a model of traumatic brain injury and parasite-induced summiting behavior.^33,42,43^

We hypothesized that increasing age and exposure to *Varroa* load would disrupt the paracellular integrity of the honeybee blood-brain barrier. Our results do not support this hypothesis, and instead suggest that paracellular integrity is robust to aging and parasitic stress (Fig. 1). The specimens collected for the *Varroa* permeability study were newly emerged, and the permeability assay was applied when they were one day old. Thus, this group also represents a younger age group than those assessed in the aging experiment. The amount of dye measured in the brains of specimens between the two experiments was similar, further suggesting that paracellular permeability is stable across lifespan and its quantification with this method repeatable across experiments. It is possible that paracellular integrity is stable over the lifespan of a honeybee and in response to stress due to the redundancy of junctional proteins within the intercellular space of SPG, though further work is needed to characterize the proteins that compose this space in honeybees.^38^

The other path a molecule can take across the blood-brain barrier is diffusion across SPG cell membranes. This transit pathway is regulated by a variety of chemoprotective mechanisms including ABC transport systems which shuttle molecules out of the barrier cells and into the hemolymph.^18^ Rho B is a small molecule dye which can diffuse passively into the blood-brain barrier layer and is a substrate for P-gp and possibly other MDR transporters which constitute a major portion of blood-brain barrier efflux systems.^35^ The concentration of brain Rho B is thus a measure in the efficiency of this major efflux mechanism at the barrier, which can be reduced by various stressors, including in honeybees.^44^ We hypothesized that aging and *Varroa* load would decrease this efficiency, thus resulting in more dye accumulation in the brains of workers with increasing age and in workers with higher mite load.

We did not find a significant difference in transcellular permeability between the age groups we examined (Fig. 2A). However, our results show a trend towards an increased transcellular permeability with age and deserve a follow up study with an increased sample size to determine whether this trend is a true effect. Increases in blood-brain barrier transcellular permeability is a observed with aging across multiple animal species including humans.^45–47^ A looming question in blood-brain barrier aging research is whether alterations in blood-brain barrier transport systems reflect a failure of function or an adaptation to the changing needs of an aging brain.^47^ The oldest honeybee workers in a colony are foragers, who engage in the collection of resources from the environment.^48^ It is possible that blood-brain barrier efflux mechanisms are downregulated during this life stage to allow for the increased uptake of endogenous molecules beneficial to the brain in a foraging context. In the modern agricultural setting however, a leaky blood-brain barrier leads to increased sensitivity to pesticides, some common types of which further downregulate the MDR efflux mechanisms in honeybees at sublethal doses.^44^ Understanding the interaction between environmental risk factors and the natural blood-brain barrier functional changes across the honeybee lifespan can inform interventions and management strategies to decrease risk factor impacts on honeybee health.^2,7^

We hypothesized that one environmental risk factor that would increase transcellular blood-brain barrier permeability is parasitism by *Varroa*. Our results support this hypothesis, specifically showing honeybees that emerged with 2-4 mites had a significantly higher brain Rho B concentration than those bees that emerged with 0-1 mites (Fig. 2B). The reproductive phase of *Varroa* occurs inside of capped brood cells, where the mites bore a hole into the cuticle of the developing brood and feed on its fat body tissue.^49^ The depletion of fat body results in organism-wide consequences such impaired development, reduced lipid synthesis, reduced protein titers, reduced hemolymph sugar levels, and impaired metabolic function.^49,50^ At the molecular level, *Varroa* exposure induces transcriptomic and proteomic alterations to immunity, oxidative stress, olfactory recognition, metabolism of sphingolipids, and RNA regulatory mechanisms.^51^ We suspect that the summed decrease in metabolic and energy resources in the specimens with a high mite load may have impacted the production of blood-brain barrier efflux transporters, resulting in increased dye accumulation.

A goal of this study was to develop and apply a blood-brain barrier permeability assay that can be widely applicable to different stress contexts. We demonstrated the efficacy of this assay to compare blood-brain barrier permeability in honeybee across two distinct lifespan stages and in honeybees exposed to varying amounts of *Varroa* infestation. While this study is the first to assess the dynamics of blood-brain barrier permeability in honeybees, further work is required to understand the specific impacts of increased blood-brain barrier permeability on measures of cognitive health in honeybees. While blood-brain barrier permeability is associated with age- and stress-related cognitive dysfunction in multiple species, it is important to understand how risk increases with degrees of increased permeability. Further, it will be interesting to probe how stress is mediated genetically and physiologically at the blood-brain barrier. For instance, a targeted study of blood-brain barrier gene and protein expression under varying amounts of *Varroa* stress will help uncover which barrier efflux proteins are most differentially expressed in response to parasitism. A similar study comparing honeybees at multiple stages of lifespan can reveal natural ontogenetic changes in blood-brain barrier function, prompting further questions about its role in supporting an aging brain.

Although age- and stress-related effects on honeybees can be subtle, it is the cumulation of risk factor impacts which are thought to underlie the devastating colony losses over the past decade.^10^ Increased blood-brain barrier permeability is often a sublethal impact, the severity of which can be rapidly assessed with this method. This assay can be particularly useful in pilot studies to identify particular concentrations or combinations of risk factors to focus on for more in depth analyses on their impact on the honeybee brain across lifespan. It may also be used as a component of monitoring studies for measuring honeybee brain health and overall healthspan over long periods of exposure to anthropogenically-altered environments.^52^

## Footnotes

1 As in all arthropods, honeybees contain hemolymph, which is analogous to blood in most respects except that it does not transport gasses. The term “blood-brain barrier” is used to avoid confusion with the existing literature on this structure across taxa.

